# Measuring single-cell density with high throughput enables dynamic profiling of immune cell and drug response from patient samples

**DOI:** 10.1101/2024.04.25.591092

**Authors:** Weida Wu, Sarah H. Ishamuddin, Thomas W. Quinn, Smitha Yerrum, Ye Zhang, Lydie L. Debaize, Pei-Lun Kao, Sarah Marie Duquette, Mark A. Murakami, Morvarid Mohseni, Kin-Hoe Chow, Teemu P. Miettinen, Keith L. Ligon, Scott R. Manalis

## Abstract

Cell density, the ratio of cell mass to volume, is an indicator of molecular crowding and therefore a fundamental determinant of cell state and function. However, existing density measurements lack the precision or throughput to quantify subtle differences in cell states, particularly in primary samples. Here we present an approach for measuring the density of 30,000 single cells per hour with a precision of 0.03% (0.0003 g/mL) by integrating fluorescence exclusion microscopy with a suspended microchannel resonator. Applying this approach to human lymphocytes, we discovered that cell density and its variation decrease as cells transition from quiescence to a proliferative state, suggesting that the level of molecular crowding decreases and becomes more regulated upon entry into the cell cycle. Using a pancreatic cancer patient-derived xenograft model, we found that the *ex vivo* density response of primary tumor cells to drug treatment can predict *in vivo* tumor growth response. Our method reveals unexpected behavior in molecular crowding during cell state transitions and suggests density as a new biomarker for functional precision medicine.

---

Cell density is determined by the cell’s dry mass composition and the fraction of cell volume occupied by water, which reflects its molecular crowding level. Although cell mass and volume can vary up to 50% in proliferating cells, cell density is tightly regulated to maintain an optimal level of molecular crowding ^1,2,3^. Environmental cues such as nutrient depletion and changes in osmolarity are known to alter molecular crowding, which impacts cellular biochemistry by altering the diffusion rate and protein conformation^1,4,5^. The coupling between crowding level and cell physiology makes cell density a key proxy for characterizing fundamental cellular processes such as proliferation, apoptosis, metabolic shifts, and differentiation^1,3^, indicating its potential as a biomarker for cellular fitness and drug response. Studies on single-cell organisms such as bacteria and yeast have reported that molecular crowding levels significantly change during cell state transitions between proliferation and dormancy, and density is thought to acutely reflect these transitions^5–8^. Whether such connections between density and proliferation exist in primary mammalian cells remains unclear, in part due to limitations in existing density measurement methods.

A major challenge for measuring cell density is achieving high sampling throughput together with high precision. Traditional gradient centrifugation methods assess cell densities on a populational level, but are slow and require a large sample size which limit their use for studying transient biological processes. Single-cell measurements reveal the heterogeneity of cell density within a population, providing insight into density regulation. Magnetic levitation methods determine the density of single cells by balancing the cell’s gravity and the buoyancy exerted by a paramagnetic medium^9,10^. Methods detecting dry-mass density (dry mass over total volume), such as quantitative phase microscopy (QPM) or Raman imaging coupled with cell volume measurements, provide alternative density measurements^11,12,13,14,15,16^. Although these methods provide subcellular resolution and single-cell tracking, experiments published thus far with mammalian cells contain tens to hundreds of single cells when measuring cell density. The suspended microchannel resonator (SMR) is a microfluidic mass sensor that has been used to measure single-cell density by measuring the buoyant mass of a cell in two types of fluids with different densities^17,18–20^. However, the throughput of this approach is limited to a few hundred cells per experiment as it requires cells to be sequentially measured in two types of fluids.

SMR and QPM devices have already achieved a throughput of tens-to-hundreds of thousands of cells per experiment^21–23^. With a streamlined volume sensing unit, the same throughput could be achieved for measuring cell density. Fluorescence exclusion microscopy (Fxm) provides a volume measurement compatible with existing SMR devices. Fxm measures the exclusion of fluorescence intensity induced by single cells that are suspended in a highly fluorescent media with cell-impermeable dye molecules. This method has been adapted to measure single-cell volumes of various model systems including bacteria, yeast, and mammalian cells^24,25^.

Here, we present a fluorescence exclusion-coupled SMR (fxSMR) platform that simultaneously measures single-cell buoyant mass and volume, allowing us to profile cell density with a throughput of over 30,000 cells per hour and a precision of 0.03% (0.0003 g/mL). We show three advances that are enabled by our high throughput and precision: i) identifying unexpected density heterogeneity, ii) revealing molecular crowding associated changes during cell state transition, and iii) validating density as a new biomarker of drug response.

## Results

### Platform design and characterization

To couple single-cell mass and volume measurements, our system is composed of an SMR cantilever with microfluidic inlets for receiving a stream of single cells and a fluorescence microscopy setup positioned at the entry of the resonator chamber (**Fig. 1a**). The fluorescence level emitted from the detection region is continuously monitored by a photomultiplier tube (PMT). To achieve the fluorescence exclusion volume measurements, cells are suspended in a fluorescent media that contains cell-impermeable dye-conjugated dextran. When there is no cell present at the detection region, the PMT detects a high fluorescence baseline from the media. As the cell passes through, the fluorescence level decreases proportionally to the volume of the cell. The raw volume signal is computed by taking the ratio of the change in fluorescence level to the baseline height (**Fig. 1b,c**).

**Fig. 1.**
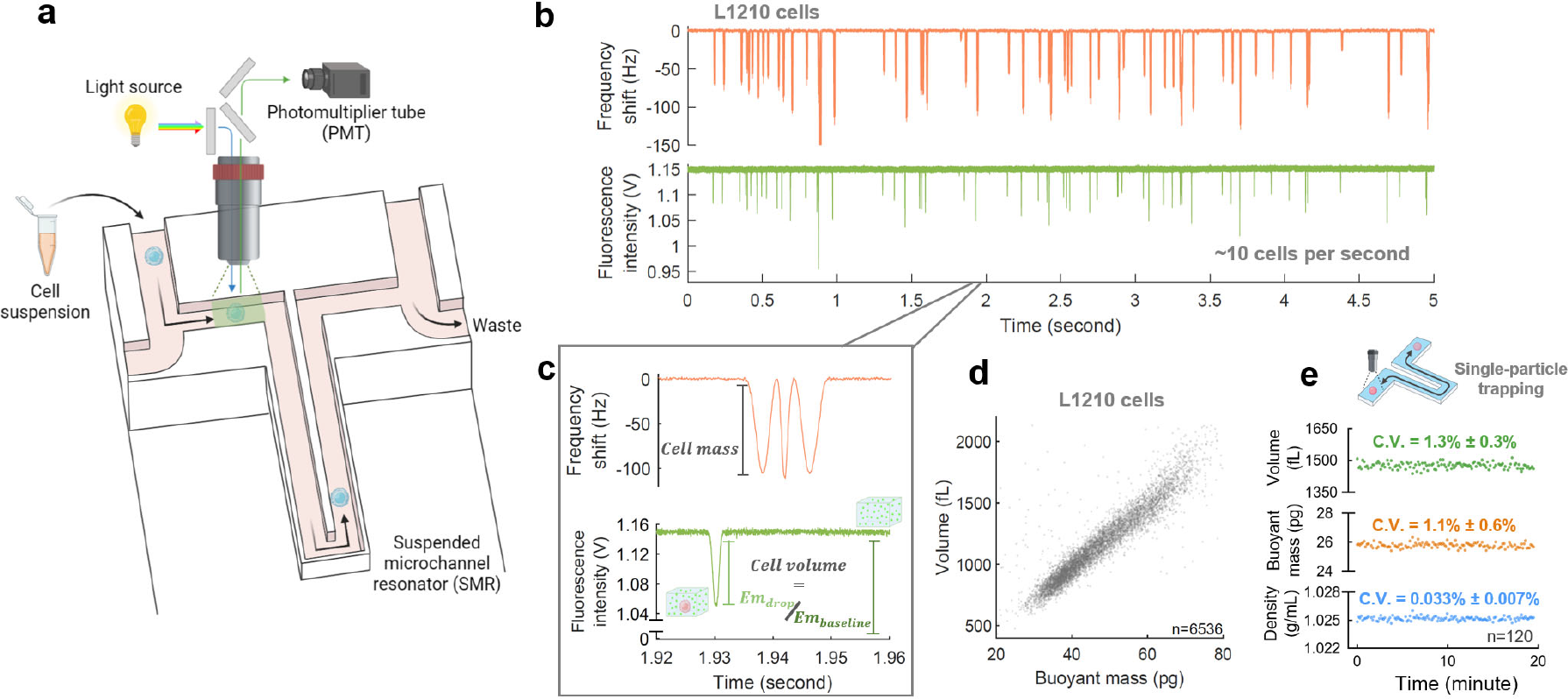
Fluorescence exclusion-coupled suspended microchannel resonator (fxSMR) enables high-throughput and high-precision single-cell density measurements. **a**, Schematic showing the system design; a fluorescence detection setup is positioned above the SMR microfluidic chip; the green shaded area indicates the fluorescence detection region and black arrows indicate the flow direction of single cells (blue). **b**, Raw signals from SMR (orange) and the photomultiplier tube (green) when measuring L1210 cells; each peak indicates the passing of a single cell. **c**, Zoomed-in image of (**b**), highlighting the shape of SMR and PMT signals for a single cell, and the calculation of cell volume from the drop in fluorescence signal (*Em*_*drop*_) from the fluorescence baseline (*Em*_*baseline*_). **d**, Scatter plot of cell mass vs volume for a population of L1210 cells as measured in ∼20 minutes; n value refers to the number of individual cells. **e**, Representative plots of volume, mass, and density of a single hydrogel microparticle that was measured repeatedly using fluidic trapping; The measurement precision (coefficient of variation, C.V.) is reported for each metric (mean ± std for 5 independently trapped particles); n value refers to the number of repeat measurements for the individual particle shown in the plots.

Cells flow through the SMR after volume measurement and the buoyant mass signal can be resolved from the change in SMR resonance frequency^17,26^. Cell density, a.k.a. buoyant density, is then computed by the following formula:

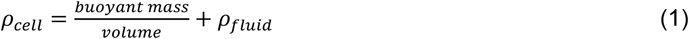

We profiled a mouse lymphocytic leukemia cell line, L1210, and simultaneously obtained single-cell buoyant mass and volume readouts with a throughput of >30,000 cells per hour (**Fig. 1b,d**). We then characterized the measurement precision by trapping single hydrogel particles which have an average diameter ∼14 μm (**Supplementary Fig. 1a**). Buoyant mass, volume, and density of the same particle were repetitively measured over a period of 20 minutes. The average coefficients of variation (C.V.) from 5 independently trapped particles were 1.3% (volume), 1.1% (buoyant mass), and 0.03% (density) (**Fig. 1e, Supplementary Fig. 1b**). Using five cell lines with median diameter ranging from ∼10 to ∼20 μm, we found a linear correlation with cell volume measured by Fxm with ground-truth volumes determined by Coulter counter (Pearson correlation coefficient R^2^ = 0.998) (**Supplementary Fig. 1d,e**). The buoyant mass measurement performance has been extensively characterized in previous studies^26–28^. These measurements demonstrate that the fxSMR platform achieves a 10 to 100-fold increase in throughput over previous approaches^9–17,20^ without compromising accuracy or precision.

### Density variation in proliferating mammalian cells

Next, we sought to examine the heterogeneity of single-cell densities within a population, as enabled by the high throughput nature of our density measurement. Most biological features with homeostatic regulatory mechanisms tend to exhibit a Gaussian distribution^29^. An exemption from this rule is the distribution of cell size, which follows a log-normal distribution due to cell growth being exponential^30,31^. A deviation from the Gaussian distribution would suggest the existence of a more complex control mechanism compared to a simple negative feedback mechanism.

We started by characterizing five suspension-grown mammalian cell lines, L1210, THP-1, BaF3, FL5.12, and s-HeLa. For statistical analysis, we gated the viable cell population (**Supplementary Note. 1**). The C.V. of cell density in all five models was below 0.6%, while the C.V. of mass and volume were considerably larger, with a range between 20-30% (**Fig. 2a**). This is consistent with previous reports on density heterogeneity in mammalian cells^1,18^. Since the density C.V. is more than 10-fold higher than our measurement precision (**Fig. 1e**,**2a**), our approach is well-suited to examine the shape of density distributions. Unexpectedly, cell densities in all five cell lines did not fit to a normal distribution (**Fig. 2b, Supplementary Fig. 2a**). We also found that a log-normal distribution did not fit well with cell density when compared to mass and volume (**Supplementary Fig. 2a-c**). The density distributions were asymmetric and better fitted by a distribution model (stable distribution) that accounts for the “heavy-tailed-ness” of the distribution (**Fig. 2b, Supplementary Fig. 2a,d**). Consistent with this, the kurtosis factors of the density distributions were all higher than 3, which is expected for a normal distribution (**Fig. 2c**). Moreover, since the hydrogel particle density measurements displayed normal distributions (**Supplementary Fig. 1c**), the higher kurtosis in cells is biological rather than a reflection of measurement bias.

**Fig. 2.**
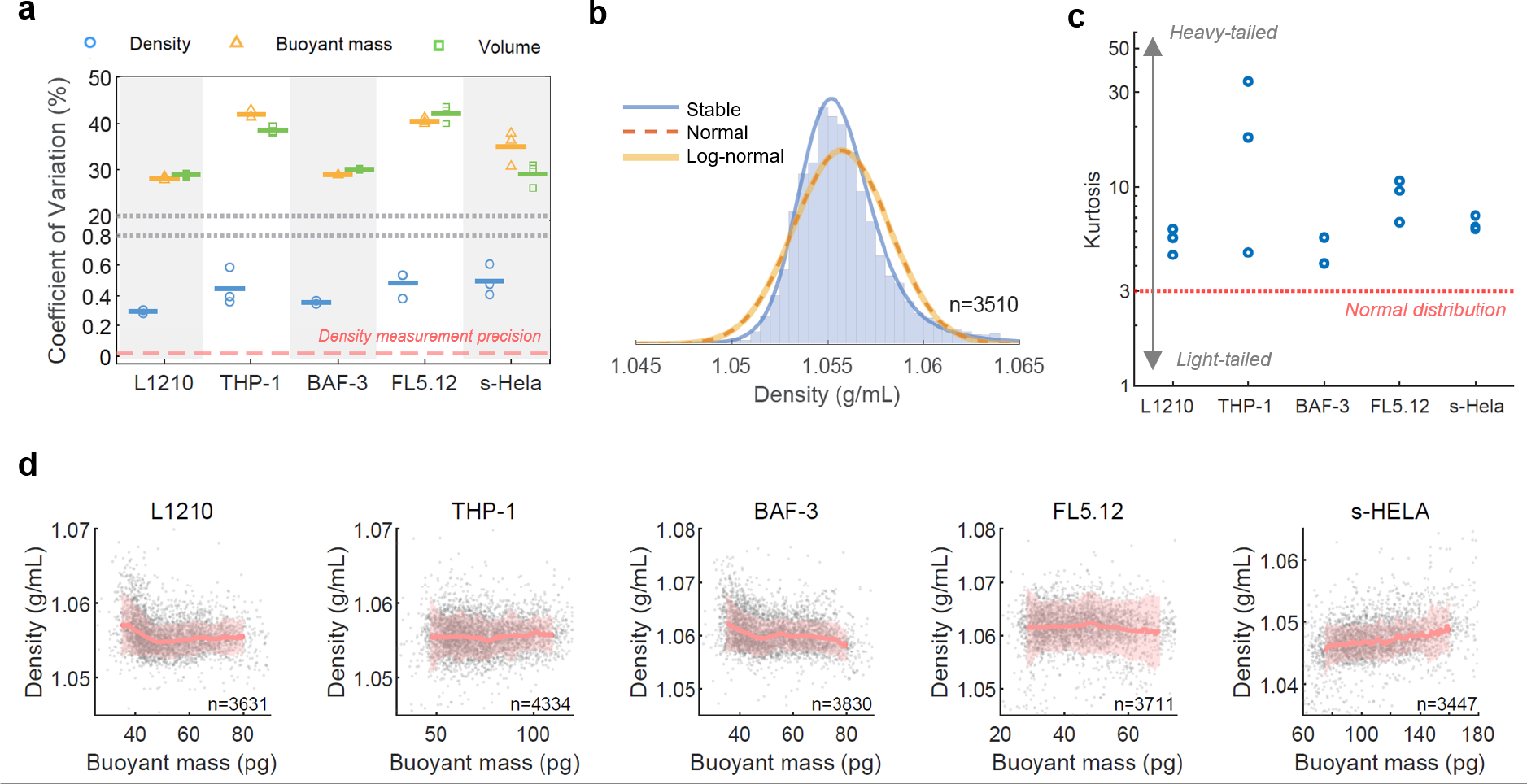
Single-cell densities have a tight and non-gaussian distribution. **a**, Coefficient of variation of density, buoyant mass, and volume for 5 cell lines; each marker represents an independent replicate, short horizontal bars denote mean; dashed red line denotes the system precision of density measurements. **b**, Density distribution of L1210 cells with lines indicating normal (red-dotted), log-normal (orange), and stable (blue) distribution fitting; n value refers to the number of individual cells. **c**, Kurtosis factor of the 5 cell lines. Each marker represents an independent replicate. **d**, Representative scatter plot of mass vs density for the 5 cell lines; Red lines and red shaded areas indicate median ± std of cell density in a moving filter along the buoyant mass axis; Gray points depict single cells, and n values refer to the number of individual cells. For (**a**), (**c**) and (**d**), n = 3 biological replicates for L1210, THP-1, FL5.12, and s-HeLa; n = 2 biological replicates for BaF3.

We next considered the cell cycle as a potential source of density variation. We used cell size as a proxy for cell cycle progression because they are tightly correlated^30,32–35^. From the five suspension cell line models, we did not observe any systematic correlation between cell mass and density, although individual cell lines show distinct features of cell cycle-dependent density (**Fig. 2d, Supplementary Fig. 3**). These results confirm that for proliferating suspension-grown mammalian cells, cell cycle progression (within interphase) does not introduce changes in cell density that would be shared between cell lines. Given prior evidence that cellular dry density doesn’t vary during the cell cycle^36^, molecular crowding level appears largely independent of the cell cycle stage within interphase. Furthermore, in every cell line, both light (<median buoyant mass) and heavy cells (>median buoyant mass) display kurtosis higher than 3, suggesting the heavy-tailed distributions are not due to a particular cell cycle stage (**Supplementary Fig. 2e**).

### Density changes during cytotoxic T-cell activation

Our results indicate that cell density does not systematically change when cells are cycling in interphase, but previous studies have revealed that cell cycle exit due to cell senescence can alter density^37^. Similarly, in single-cell organisms, cells can enter a quiescence state where the cells display higher molecular crowding (i.e. higher density) and smaller cell size^5–7^. We therefore examined if density homeostasis is specific to cycling cells (**Fig. 3a**). To study this, we focused on models where we can compare cell density in quiescence and proliferative states.

**Fig. 3.**
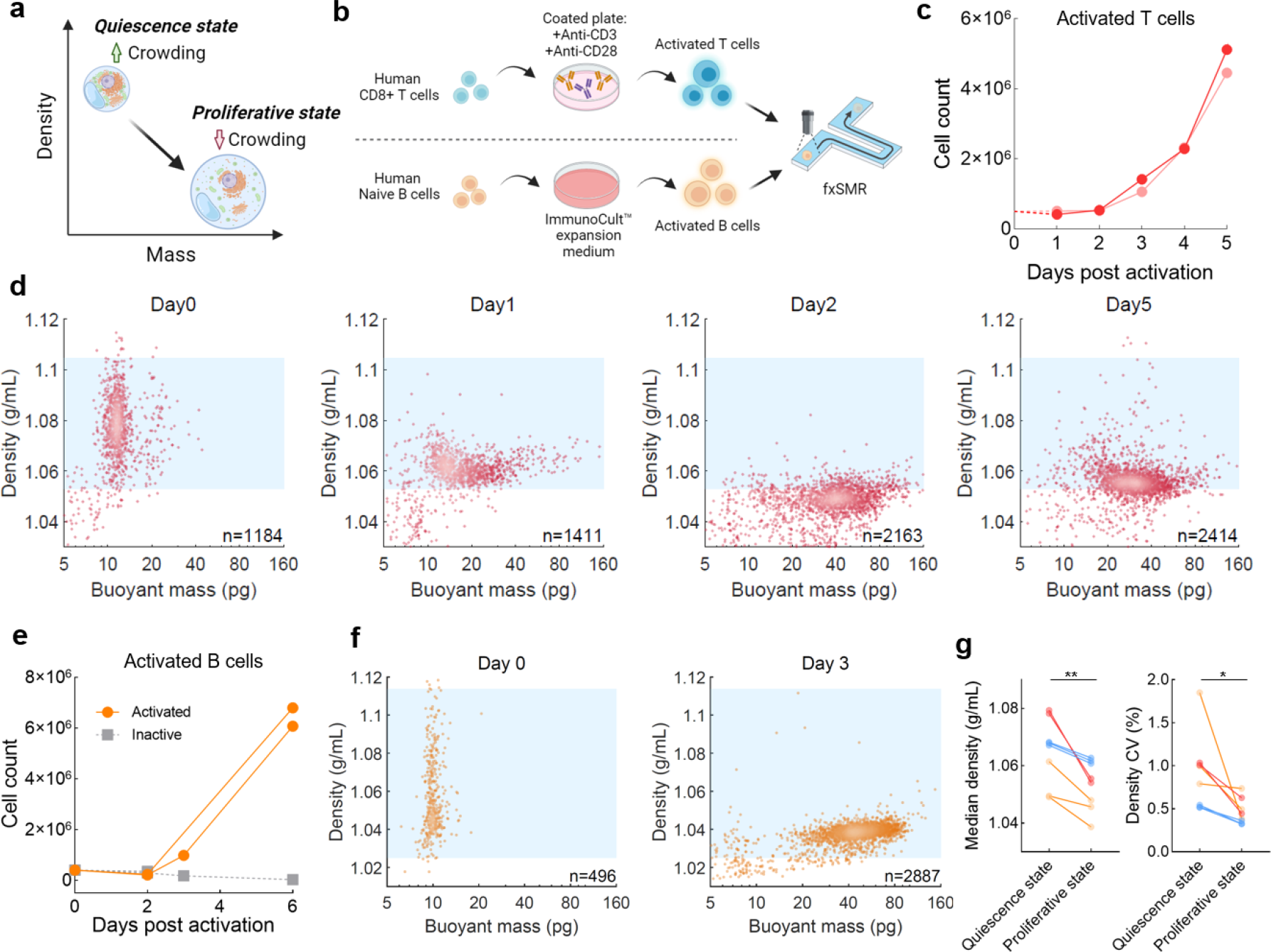
Density profiling of human lymphocyte activation shows crowding transitions between quiescence and proliferative states. **a**, A qualitative model of biophysical changes associated with the transition between quiescence and proliferative states. **b**, Schematics showing the activation process of human CD3+/CD8+ T cell and human naive B cell and subsequent profiling by fxSMR. **c**, Cell count vs time (days post activation) for T cells obtained from two donors (red and pink); Dotted line denotes expected seeding density. **d**. Scatter plots of mass vs density showing the T cell dynamics post activation for donor 1; Blue areas indicate the density range of quiescent T cells at day 0, with upper and lower bounds indicate the 1st and 99th percentile of the density distribution; C.V. of density at day 0, 1, 2 and 5 are 1.008%, 0.453%, 0.392%, 0.627% accordingly; n values refer to the number of individual cells. **e**, Representative cell count vs time (days post activation) for naive B cells obtained from two donors. The B cells were either activated (solid orange lines) or kept as inactive naïve B cells (dotted grey line). **f**. Representative scatter plots of mass vs density showing B cells at day 0 and 3 post activation; Blue areas indicate the density range of quiescent B cells similar to (**d**). C.V. of density at day 0 and 3 are 1.850% and 0.384% accordingly; n values refer to the number of individual cells. **g**. Median cell density and the C.V. of density for quiescent and proliferating lymphocytes. Two-tailed paired t tests yield p-value = 0.0049 for median density, and p-value = 0.0269 for density C.V.; Red indicates T cells (n = 2 biological replicates); Orange indicates B cells (n = 3 biological replicates); Blue indicates murine pro-B lymphocytic cell line FL5.12 (n = 3 biological replicates).

First, we studied human lymphocytes from peripheral blood that circulate as quiescent cells and can readily become activated and proliferating effector cells after encountering external stimuli. We performed daily measurements of CD8+ T cells from two human donors after co-stimulation with anti-CD3/CD28 (**Fig. 3b,c**). During the first two days post-activation, the density C.V. decreases from ∼1% to ∼0.4% suggesting a stronger density regulation as T cells start to proliferate (Chi-square variance tests reported p-values < 0.00001 for both donors) (**Supplementary Fig. 4g**). In the same timeframe, T cells increased their size substantially while decreasing average density from ∼1.08 to ∼1.05 g/mL (**Fig. 3d,g, Supplementary Fig. 4a**). We then sought to determine if this decrease in average cell density reflects changes in molecular crowding. As a proxy for molecular crowding, we measured the fraction of osmotically active water content over total cell volume by applying the Boyle Van’t Hoff relation, where the volume of a cell is inversely proportional to the external osmolality^38–40^.We measured the active water content of T cells by profiling the volumes of each sample under two different osmolarity conditions (**Supplementary Fig. 4b**). We found that T cells significantly increased their water content from ∼63% to ∼80% of total cell volume within the first two days of activation (**Supplementary Fig. 4c,d**), suggesting a lower crowding level before cells start to divide. The relationship between cell density and intracellular molecular concentrations is discussed in more detail in **Supplementary Note. 2**.

We confirmed our findings by studying different cellular models of cell quiescence and proliferation. Similar trends in cell density and size were observed in activated and proliferative human naïve B cells (**Fig. 3b,e-g, Supplementary Fig. 4e-g**). In addition, we studied a pro-B lymphocytic murine cell line FL5.12, which is interleukin-3 (IL-3)-dependent and exits the cell cycle following IL3-depletion^41^. As with the human lymphocytes, we observed that quiescent FL5.12 cells have a higher density due to increased crowding and higher density C.V. when compared to their proliferative state (**Fig. 3g, Supplementary Fig. 5**). We performed small particle trapping experiments to validate that the observed differences in density C.V. were predominantly driven by biological variability rather than increased measurement noise in small cells (**Supplementary Note. 3**). Overall, our results reveal that lymphocytes maintain lower cell density and tighter density homeostasis when proliferating. This suggests that density regulation is coupled to the molecular machinery responsible for cell growth and/or proliferation.

### Density as a biomarker for drug response

Since changes in cell density can reveal state transitions related to cell proliferation, we sought to determine if it could be used as a biomarker for assessing *ex vivo* treatment response of cancer cells. A major goal for precision cancer medicine is to match each patient with the most effective drug treatment. Functional precision medicine (FPM) approaches, which involve drug testing performed directly on patient tumor cells, have emerged in recent years to help select the optimal drug treatment at the time of diagnosis or relapse^42^. In these assays, live cells isolated from patient tumors are treated with a panel of candidate drugs *ex vivo*, and drug responses are assessed for their ability to predict patient outcomes^43^. Proliferation-based assays for drug response work well in cell line models that have adapted to thrive in *ex vivo* conditions. However, primary cancer cells often do not proliferate or require specific culture conditions to stimulate proliferation *ex vivo*, thereby increasing risk of phenotypic drift^44^. Consequently, there is a need for predictive FPM biomarkers that minimize phenotypic drift by assessing *ex vivo* treatment response at short timescales where cells are not proliferating. A number of studies have shown that cell mass can function as a predictive biomarker with 1-2 day turnaround times^21,22,45,46^. We hypothesized that cell density response could also function as a hyper-acute predictive biomarker even in the absence of proliferation, given that increases in crowding level (i.e. cytoplasmic condensation) have long been regarded as an early indicator of apoptosis^18,47,48^.

To validate the density biomarker, we used a patient-derived xenograft (PDX) model as a source of cells to benchmark the density response against previously established FPM assays of proliferation and cell mass measurements (**Fig. 4a**). The pancreatic ductal adenocarcinoma (PDAC) model, NIBRX-1362, harbors the KRAS G12D mutation and has known *in vivo* drug response profiles from a previous study^49^ (**Fig. 4b**). Repeat testing confirmed the model exhibited *in vivo* sensitivity to trametinib, a MEK pathway inhibitor, and only mild response to gemcitabine, a chemotherapy. We next isolated dissociated single cells derived from the untreated PDX tumors and assessed proliferation response using CellTiter-Glo at 6 days after *ex vivo* drug treatment. Using this approach, we noted a marked *ex vivo* tumor cell response to trametinib and a less effective response to gemcitabine (IC50 = 0.172 nM and 8.561 nM respectively) (**Fig. 4c**) thereby validating that the *ex vivo* response agreed with the *in vivo* results at an “acute” 6-day time frame.

**Fig. 4.**
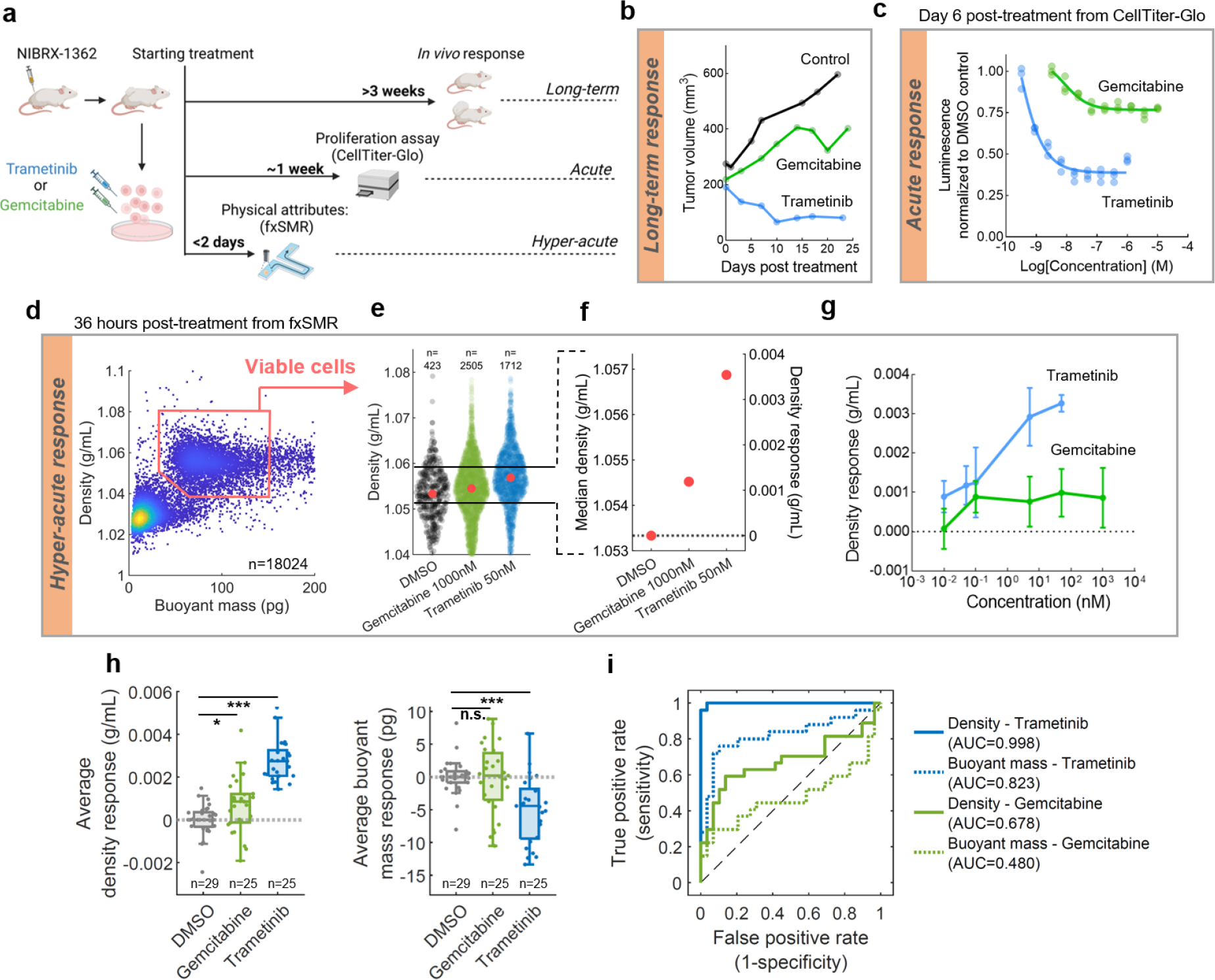
Density demonstrates robustness as a hyper-acute biomarker for predicting long-term *in vivo* drug response. **a**, Schematics showing the paradigm of functional biomarkers for predicting long-term *in vivo* treatment response. NIBRX-1362 is a pancreatic ductal adenocarcinoma (PDAC) patient-derived xenograft (PDX) model. Disassociated singles cells were used for proliferation assays and fxSMR measurements of cell density and mass. **b**, *In vivo* treatment response measured by tumor volume data (1 mouse per condition) (ref.^49^). **c**, CellTiter-Glo dose response at day 6 after treatment, with 3 biological replicates per drug concentration; Slopes represent dose response fitting; IC_50_ = 0.1724 nM for trametinib and 8.561 nM for gemcitabine. **d**, Representative scatter plot of mass vs density at 36 hours post-treatment, after pooling together single-cell data of DMSO, 1000 nM gemcitabine, and 50 nM Trametinib treatments; Red box shows gating of viable cells; n value refers to the number of individual cells. **e**, Viable single-cell densities are plotted for indicated conditions. Red dots denote median densities, n values refer to the number of individual cells. **f**, Calculation of density response for each treated condition using median densities; The dotted line indicates where density response = 0, i.e. median density equal to the DMSO control. **g**, 36-hour density responses for gemcitabine and Trametinib at multiple concentrations; EC_50_ for trametinib treatment is 0.017 nM, and cannot be stably fitted for gemcitabine treatment; Data indicates mean ± SEM; n = 3-6 biological replicates. **h**, The average density (left) and buoyant mass (right) responses of DMSO, gemcitabine and trametinib treatments. Based on the dose-response result in (**g**), high drug doses (≥10 nM for both gemcitabine and trametinib treatments) were grouped for each treatment time. From unpaired parametric t tests, the average density responses have p-value = 1.391×10^-16^ for trametinib vs DMSO treatment; p-value = 0.018 for gemcitabine vs DMSO treatment. Average buoyant mass responses have p-value = 2.139×10^-5^ for trametinib vs DMSO; p-value = 0.808 for gemcitabine vs DMSO treatment. **i**, Receiver operating characteristic (ROC) analysis of density (solid lines) and buoyant mass (dotted lines) responses using data from (**h**); the plot indicates the robustness of a binary prediction of *in vivo* treatment response against DMSO control (prediction on trametinib treatment shown as blue, and gemcitabine shown as green). True-positive rate indicates the occurrence rate of the response giving the correct prediction (over the threshold) when the PDX model is sensitive to the *in vivo* treatment; False-positive rate indicates the rate of the response giving the incorrect prediction (under the threshold) when the PDX model is sensitive to the *in vivo* treatment. Area under the curve (AUC) scores denote the predictive powers of density and mass response.

To determine whether single-cell density can achieve the same conclusion but at a hyper-acute timescale, we profiled single-cell density and mass using fxSMR at 24 and/or 36 hours post drug treatment, using similar conditions to the CellTiter-Glo assay. We first applied a gating based on density and buoyant mass to select viable cells (**Fig. 4d, Supplementary Note. 1**), and then for every drug-treated sample, we subtracted the median density of the sample from the median density of the control (**Fig. 4 e,f**) to obtain a density response for each treatment. A dose-response analysis demonstrated that both trametinib and gemcitabine resulted in density response when compared to the DMSO treatment (n=9 separate experiments and PDX bearing mice) (**Fig. 4g**). At high concentrations (≥10 nM), trametinib treatments displayed a greater and more significant density response compared to gemcitabine (median density response = 0.0027 g/mL and 0.0009 g/mL, accordingly), which agreed with the long-term *in vivo* response and the CellTiter-Glo results (**Fig. 4h**).

Furthermore, when benchmarking the ability to distinguish between treatment responses, we found that buoyant mass detected a significant response from trametinib treatment, but could not resolve a significant gemcitabine response (**Fig. 4h**). To further test whether the density response is more likely to align with the long-term *in vivo* response, we performed a ROC analysis which showed that, for both treatments, density has a stronger predictive power than mass (for trametinib, density AUC = 0.998 and mass AUC = 0.823; for gemcitabine, density AUC = 0.678 and mass AUC = 0.480) (**Fig. 4i**). Since fxSMR readouts were collected from a cohort of 9 replicates of the PDX model, the strong AUC score for predicting trametinib response also suggests that *ex vivo* cell density response is highly robust in resolving the long-term *in vivo* response of an effective treatment observed in this PDX model.

## Discussion

We have shown that fxSMR can precisely measure single-cell density with 10-100x higher throughput than existing methods. With our approach, we discovered that cell density and molecular crowding differ between proliferating and quiescent lymphocytes. This discovery bears resemblance to similar observations in unicellular organisms, where the increased molecular crowding in the quiescence state has been associated with slower signaling and lowered metabolic activity^4,5,50^. It is possible that the high cell density we observed in quiescent mammalian cells has a similar role where non-proliferating and largely inactive cells conserve energy. More broadly, our results suggest a conserved mechanism that couples the regulation of cell density and molecular crowding to the regulation of cell state between quiescence and proliferation. It is also worth noting that a seemingly small change in cell density (<5%) can reflect significant changes in molecular concentrations inside the cell, with potential implications to phase transitions and enzymatic reaction rates (**Supplementary Note. 2**).

Our results also reveal insights into the regulation of cell density homeostasis. We discovered that proliferating mammalian cells display less cell-to-cell density variability than quiescent cells, indicating that the strength of density homeostasis is dependent on the cell cycle machinery responsible for cycling cell state. However, cells in early and late interphase display similar density homeostasis, ruling out density regulation by G1, S or G2 cell cycle stage specific mechanism. Furthermore, our work revealed that cell density distributions are non-Gaussian in proliferating cells, thus narrowing down the space of theoretical models that could explain how density homeostasis is maintained.

In addition to its potential for exploration of the biology of density homeostasis, our approach may provide a much-needed method for functional precision medicine in patients. The ability of high-throughput single-cell density measurements to generate drug response data in the hyper-acute non-proliferative window is a unique capability when compared to other functional precision medicine approaches. For example, organoid testing typically requires longer time periods (e.g. weeks) for *ex vivo* expansion to generate a sufficient number of cells for assessments.

There are two important limitations of our drug response study. First, since we have not measured density response across multiple patient models, the broader predictive capability across heterogeneous cancers will require future studies. Second, since we have only measured the density response from two drugs, the extent to which density response will generalize across other drugs with a wide range of mechanisms remains unknown. However, previous studies with cell lines have shown that drugs with various mechanisms of action can uniformly perturb cell density^18,18,19^. With our high throughput approach, we believe that many drugs can now be more readily profiled on patient samples within a hyper-acute time window, thereby enabling clinical studies for assessing the effectiveness of density response in guiding patient treatment.

## Methods

### System setup

SMR devices were designed as previously described^51^. The fabrication was carried out at CEA-LETI in Grenoble, France, with procedures documented in previous studies^17,26^. An SMR device was actuated by a piezo-ceramic plated underneath the chip, which allows the suspended cantilever to vibrate at the resonant frequency of its second vibrational mode. Vibrational frequencies were measured by piezo resistors at the base of the cantilever. A closed-loop feedback system was applied to ensure consistent actuation at the resonant frequency, with a pre-defined delay time between the piezo resistor readout signal and the actuator driving signal. The driving signal was amplified to achieve high oscillation amplitude as well as low-frequency noise.

The Optical setup was built with an epi-fluorescence microscope (Nikon LV-UEPI2), using similar designs as previously described^52^. To reduce the noise in fluorescence measurements resulting from mechanical instabilities, two additional optical posts (Thorlabs) were installed to better support the optical pedestal (Thorlabs) between the microscope and the lower breadboard that holds the SMR device. Fluorescence excitation was provided by a laser-LED multi-band illuminator (Lumencor SPECTRA Light engine). A 50X/0.55 objective lens (Nikon-CFI, LU Plan ELWD WD 10.1mm) was used, and the emission collection area was defined by two orthogonally-placed adjustable mechanical slits (Thorlabs VA100/M). The emission light was collected using a CMOS (complementary metal-oxide semiconductor) camera (FLIR, BFS-U3-13Y3M-C), and photomultiplier tubes (Hamamatsu, H10722-20). A 10/90 beam splitter was positioned in front of the camera to direct 10% of all emission light to the camera. For each photomultiplier tube, the light path consisted of a dichroic mirror (Semrock), an emission filter (Semrock), and a convex lens (Thorlabs LA1027). The emission light was separately collected into 5 PMTs with the following emission filter ranges(nm): 438/24, 515/30, 595/31, 678/70, and 809/81 nm. Volume measurements from FITC-dextran dye molecules were conducted with ∼500 mW excitation light (475/28 nm) and the emission light was captured within the 515/30 nm band. The exact optics configurations were shown in **Supplementary Fig. 6a**. For communications with the PMTs, reference voltages were set by analog output modules (National Instruments NI-9263) and the output signals were collected by voltage modules (National Instruments NI-9215) were used.

### System operation

SMR devices have four fluidic inlets that are connected to the sample, buffer, and waste reservoirs with 0.007-inch-inner-diameter fluorinated ethylene propylene tubing (IDEX Health & Science). Pressurized house air was used to drive the fluidics. Flow directions were controlled by electronic pressure regulators and solenoid valves, through a custom software in LabView2020. The typical flow rate was around ∼10 nL/s, as estimated by the time for an average particle to travel through the SMR cantilever.

For a typical single-cell density measurement, cells were resuspended to a final concentration of ∼1 million cells/mL in the fluorescent buffer. The buffer was made by dissolving 2000 kDa Fluorescein isothiocyanate– dextran (Sigma, FD2000S) in PBS or cell culture media to a final concentration of 10 mg/mL. Each sample was typically measured for 15-20 minutes. During the run, the sample reservoir was stored on ice to minimize endo/exocytosis. After each measurement, the SMR cantilever was flushed with 50% bleach in water and subsequently with PBS to minimize any accumulation of debris or air bubbles along the channel wall. To ensure consistent volume measurement across different samples, we adjusted the positioning of rectangular slits at the start of each experiment to define the emission collection area (**Supplementary Fig. 6b**,**c**). Given that channel height was fixed, the slits configuration determined the total fluorescence excitation volume, which was used to calculate cell volume. Slits configuration was usually set to 20 μm (W) by 40 μm (L) for a typical sample. The width was further adjusted for samples with very small average cell sizes (naive T cells), and large average cell sizes (s-HeLa). Data acquisition was enabled by custom measurement software in LabView2020 for both SMR and optical readouts. SMR data were acquired at a data rate of ∼20 kHz and light-intensity readouts from PMTs were acquired at a rate of ∼50 kHz. Subsequent fxSMR data analysis is described in **Supplementary Note 4**.

### Calibration of volume and buoyant mass measurement

Calibration of the raw volume signal from fxSMR to the standard unit (femtoliter) was done with the L1210 cell line. Before each experiment, we first measured a cell line population in PBS suspension using a coulter counter (Beckman Coulter Multisizer 4, 100 μm aperture) to obtain a cell volume distribution with the standard unit. Then, we measured the cells using fxSMR and acquired a distribution of their raw volume signals. From these two distributions, we first took the median of each measurement and calculated their ratio as an estimated calibration factor (Volume-in-fL/Volume-in-a.u.). Then we will refine the calibration accuracy by looping through a range of potential calibration factors (± 3 a.u. around the starting calibration factor derived by medians) to minimize the difference between the two distributions (the summed difference in probability density between each percentile in the distribution). This optimization step led to a final calibration factor. SMR frequency peaks were calibrated (hertz to picogram) with 8 μm polystyrene beads with a known density of 1.05 g/mL (Thermo Fisher, Duke Standards). Calibration calculations for both cell volume and buoyant mass were conducted using MATLAB.

### Accuracy and precision characterization

We characterized the accuracy of fxSMR volume measurement by benchmarking against coulter counter volume measurement using five different cell lines, with median cell size ranges from ∼1000 to ∼5000 fL. The volume of each cell line sample was first determined by a Coulter Counter (Beckman) and subsequently measured by fxSMR. From the single cell volume distributions measured by the two platforms, median value of each sample was computed and used to calculate correlation score between fxSMR measurements and ground-truth cell volumes. (**Supplementary Fig. 1c,d**). The precision of fxSMR was determined by repeatedly measuring the same hydrogel particle (deformable poly-acrylamide-co-acrylic acid microparticle), donated by Morgan Huse lab at the Sloan Kettering Institute. Each particle was pushed forward and backward through the cantilever at the rate of ∼30 seconds per measurement for a duration of ∼20 minutes. The trapping mechanism was achieved by oscillating the pressure setting between the left and right side of the cantilever to generate forward and backward pressure gradients. Real-time SMR frequency analysis was used to recognize the passage of a particle and the pressure setting was then switched after a pre-defined period of time (50-200 ms) after each passage. Fluidic controls were carried out through a custom software in LabView2020. The ground-truth particle size was determined from 60x brightfield images taken using an imaging flow cytometer (Amnis ImageStream, Cytek). Particle areas were calculated using automated feature identification using the IDEAS image analysis software. Particle volumes were subsequently calculated using GraphPad Prism.

### Statistics

Statistical analysis was conducted using MATLAB and GraphPad Prism. Statistical significance between groups or different experimental conditions was determined using unpaired parametric t test in **Fig. 4**, and paired parametric t test in **Fig.3** and **Supplementary Fig. 5**. Chi-square variance test for **Supplementary Fig. 4g** was performed with vartest() MATLAB function. Distribution fitting over single cell data in **Fig. 2** was conducted with *fitdist()* MATLAB function with specification on distribution type such as normal, log-normal or stable. Kurtosis and skewness factor of single cell distributions were calculated by *kutosis()* and *assness()* MATLAB functions. One-sample Kolmogorov-Smirnov tests in **Supplementary Fig. 2a-c** were conducted by first fitting the data with the intended distribution type using *fitdist()*, and then *kstest()* MATLAB function was performed to test the data against the cumulative distribution function of the fitted model. Bootstrap analysis for **Fig. 2** and **Supplementary Fig. 2a-c** was carried out with the *bootstrp()* MATLAB function with an 1000 iterations. ROC analysis in **Fig. 4** was conducted with the *perfcurve()* MATLAB function. A binary sensitivity label was given as the outcome variable, with trametinib-treated or gemcitabine-treated samples as sensitive and DMSO-treated conditions as non-sensitive.

### Water content measurement by osmotic shocks

Osmotically active water content was measured by volume exclusion on fxSMR. We measured aliquots from the same sample population that were resuspended in different osmolarities. Isotonic buffer is the cell culture media containing 5mg/mL FITC-dextran, which has an estimated osmolarity of 300 mOsm. The 600 mOsm hyperosmolarity buffer was made by first diluting NaCl stock solution (Sigma S5150) with water to a final concentration of 450mM NaCl in water, and then mixing the NaCl buffer with PBS that contains 10 mg/mL FITC-dextran, with a 1:1 mixing ratio. After fxSMR volume measurements on the two buffer conditions, the median cell volume was calculated from each single-cell dataset. A linear regression fitting was conducted using median cell volumes as y values and the corresponding [1/osmolarity] as x values. The y-intercept of the fitted slope was determined as the osmotically inactive volume, and osmotically active water content is determined by subtracting the inactive volume from the median cell volume of the isotonic measurement (**Supplementary Fig. 4b)**.

### Cell culture

L1210, THP-1, BaF3, FL5.12, s-HeLa, and HL60 cells were cultured in RPMI (Invitrogen). Patu-8902 cells were cultured in DMEM (Gibco). For all cells, the media was supplemented with 10% FBS (Sigma-Aldrich), 1 mM Na pyruvate (Invitrogen), 10 mM HEPES (Invitrogen), and antibiotics (Invitrogen). The FL5.12 cell culture media was also supplemented with 100 ng/ml IL-3 (R&D Systems). L1210 cells were obtained from ATCC, THP-1 cells were graciously donated by the Chen lab at MIT, BaF3 cells were obtained from RIKEN BioResource Center, FL5.12 cells were previously donated by the Vander Heiden lab at MIT. s-HeLa cells were previously donated by the Elias lab at Brigham and Women’s Hospital. HL60 cells were previously donated by Thiam lab at Stanford. Patu-8902 cells were previously donated by the Hahn lab at the Broad Institute. All experiments with cell lines were carried out with exponentially growing cells at a confluency of 300,000 – 600,000 cells/mL. All cell lines were tested for mycoplasma and no mycoplasma was detected. For FL5.12 IL-3 depletion, cells were first grown to confluency at 1 million cells/mL, and subsequently washed three times in RPMI media without IL-3 and resuspended in RPMI media without IL-3 at a concentration of 500,000 cells/mL.

For CD8+ T cell activation, apheresis leukoreduction collars from anonymous healthy platelet donors were obtained from the Brigham and Women’s Hospital Specimen Bank under an Institutional Review Board– exempt protocol. Human peripheral blood mononuclear cells (PBMCs) were isolated via density gradient centrifugation (Lymphoprep, StemCell Technologies Inc, cat#07801). PBMCs were resuspended in cryopreservation media (90% FBS + 10% DMSO) and frozen down. For T cell activation, 24 well plates were precoated with anti-human CD3 antibodies (0.5 ug/mL, BioXCell, cat#BE0001-2) and anti-human CD28 antibodies (5 ug/mL, BioXCell, cat#BE0248) for 24 hours at 4°C. PBMC samples were thawed on the day of activation and T cells were purified via the EasySep™ Human CD8+ T Cell Enrichment Kit (StemCell Technologies Inc, cat#19053). Isolated T cells were seeded in the CD3/CD28 precoated well plates at a concentration of 500,000 cells/mL with 1 mL/well in ImmunoCult™-XF T Cell Expansion Medium (StemCell Technologies Inc, cat#10981) supplemented with 10 ng/mL recombinant IL-2 (StemCell Technologies Inc, cat#78036.1). T cells were removed from CD3/CD28 plate coating after 48 hours of incubation and moved to uncoated wells. For the remainder of the culture, cells were passaged when the concentration reached over 1 million cells/mL to a new concentration of 500,000 cells/mL.

For B cell activation, apheresis leukoreduction collars from three anonymous healthy platelet donors were obtained from the Brigham and Women’s Hospital Specimen Bank under an Institutional Review Board– exempt protocol. PBMCs were isolated with Ficoll-Paque Plus (Thermo Fisher Scientific, # 45001749) using the manufacturer’s recommended protocol. The PBMC layer was isolated, subjected to ACK lysis (Life Technologies, # A1049201), and washed with PBS. Naïve B cells were isolated using EasySep™ Human Naïve B Cell Isolation Kit (Stem Cell, #17254) according to the Manufacturer’s protocol. The Naïve B cells were seeded at 100,000-250,000 cells/mL in a 6-well plate and cultivated in the ImmunoCult™ Human B Cell Expansion Kit (Stem Cell, #100-0645). The plating density was adjusted to 100,000 cells/mL every 2-4 days and the viability was assessed by Trypan blue. Unstimulated naïve B cells were cultivated in RPMI supplemented with 10% FBS and antibiotics. The immunophenotype was confirmed by flow cytometry. The cells were stained in Brilliant Stain Buffer (BD Biosciences, # 566349) with the following dyes and antibodies diluted at their predetermined optimal concentrations: Zombie Aqua (BioLegend, #423102), anti-human CD19 (BD Biosciences, #564456), anti-human IgD (BD Biosciences, #561314), anti-human CD27 (BD Biosciences, #562513), anti-human CD86 (BD Biosciences, #555665), anti-human HLA-DR (BD Biosciences, #560743), anti-human CD24 (BD Biosciences, #563401) and anti-human CD38 (BD Biosciences, #555462). The cells were then washed with PBS supplemented with 2% FBS and 0.2% EDTA. Stained cells were analyzed on a BD LSR Fortessa flow cytometer (Dana-Farber Cancer Institute Flow Cytometry core), and data were analyzed using FlowJo v10 software.

### PDX model development and *ex vivo* drug testing

The NIBRX-1362 PDAC PDX model was previously established and characterized^49^ and was obtained from the Dana-Farber Cancer Institute (DFCI) Center for Patient Derived Models which is a distributor of the model (https://www.dana-farber.org/research/departments-centers-and-labs/integrative-research-centers/center-for-patient-derived-models/ordering-patient-derived-models/). The cryopreserved PDX seeds were thawed and rinsed with DMEM before being implanted in athymic nude mice (Taconic, NCRNU-F) in the presence of matrigel (Corning). When the tumor reached a palpable size, tumor volume was measured and monitored using a digital caliper. Tumors were harvested when approaching 2000 mm^3^ and were serially passaged from harvested tumor seeds. The identity of the NIBRX-1362 PDX model used in this study was confirmed via STR fingerprinting and key gene mutations (KRAS G12D) were verified via exome sequencing.

To perform the acute drug sensitivity assays, freshly harvested PDX tumors were aseptically explanted from the flank, and processed to remove necrotic tumor regions and mechanically dissociated into single cells using the Miltenyi Tumor Dissociation kit (Miltenyi, 130-095-929) and the GentleMax dissociator according to the manufacturer’s instruction. The dissociated cells were filtered using a 0.2 μm strainer and subjected to mouse cell removal using a mouse cell depletion kit (Miltenyi, 130-104-694). The mouse-depleted tumor cells of NIBRX-1362 were plated at 1250 cells per well with 40 μL of DMEM+2%FBS (Gibco, 11965092; Sigma, F2442) in 384-well opaque plates (Corning, 3570) for acute drug sensitivity measurement via CTG and plated at 75k cells per well with 3 mL of DMEM+2%FBS in 6-well ultra-low attachment plates (Corning, 3471) for fxSMR assays.

For acute drug sensitivity measurement using the CTG assay, the freshly plated PDX cells were incubated overnight before treatment with an 8-concentration range of gemcitabine (10 μM - 3.25 nM, SelleckChem S1714) or trametinib (1 μM - 0.325 nM, SelleckChem S2673) using an automated drug dispenser (Tecan D300e Digital Dispenser). After 6 days of drug treatment, cells were examined using CellTiter-Glo 2.0 cell viability assay (Promega G9241) according to manufacturer instruction. Data was analyzed using GraphPad Prism.

## Acknowledgements

We thank M. Huse (Sloan Kettering Institute) for donating DAAM-particles, J. Chen (Koch Institute) for donating THP-1 cells, M. Vander Heiden (Koch Institute) for donating FL5.12 cells, K. Elias (Brigham and Women’s Hospital) for donating s-HeLa cells, H. Thiam (Stanford) for donating HL60 cells, and B. Hahn (Broad Institute) for donating Patu-8902 cells. We thank the Koch Institute’s Robert A. Swanson (1969) Biotechnology Center for technical support, specifically The Swanson Biotechnology Center Flow Cytometry Facility. Schematics presented in this work were created with BioRender.com. S.R.M. discloses support for the research described in this study from Paul G. Allen Frontiers Group, Virginia and D.K. Ludwig Fund for Cancer Research, MIT Center for Precision Cancer Medicine and Stand Up to Cancer (SU2C) Convergence Program 3.1416. K.L.L. and S.R.M. disclose support for the research described in this study from Bristol Myers Squibb. This work was also supported in part by the Koch Institute Support (core) Grant P30-CA014051 from the National Cancer Institute

## Author contributions

W.W., T.P.M., K.L.L., and S.R.M. conceptualized the study. W.W. and S.H.I. designed and built the measurement device with assistance from T.P.M and Y.Z. For accuracy and precision characterizations, T.P.M carried out cell culture work, and W.W. and S.H.I. performed fxSMR measurement and data analysis. For the lymphocyte study, S.M.D. and L.D. carried out the cell culture work, and W.W. and Y.Z. carried out fxSMR measurement and data analysis. For the PDX study, T.W.Q., S.Y., P.L.K. and K.H.C. carried out all *in vivo*, CTG and cell culture work, and W.W. and S.H.I. performed fxSMR measurement and data analysis. W.W. wrote the manuscript with contributions from M.M., M.A.M., T.P.M., K.L.L. and S.R.M. All authors reviewed and approved the manuscript.

