## Supplementary Information for "Measuring single-cell density with high throughput enables dynamic profiling of immune cell and drug response from patient samples"

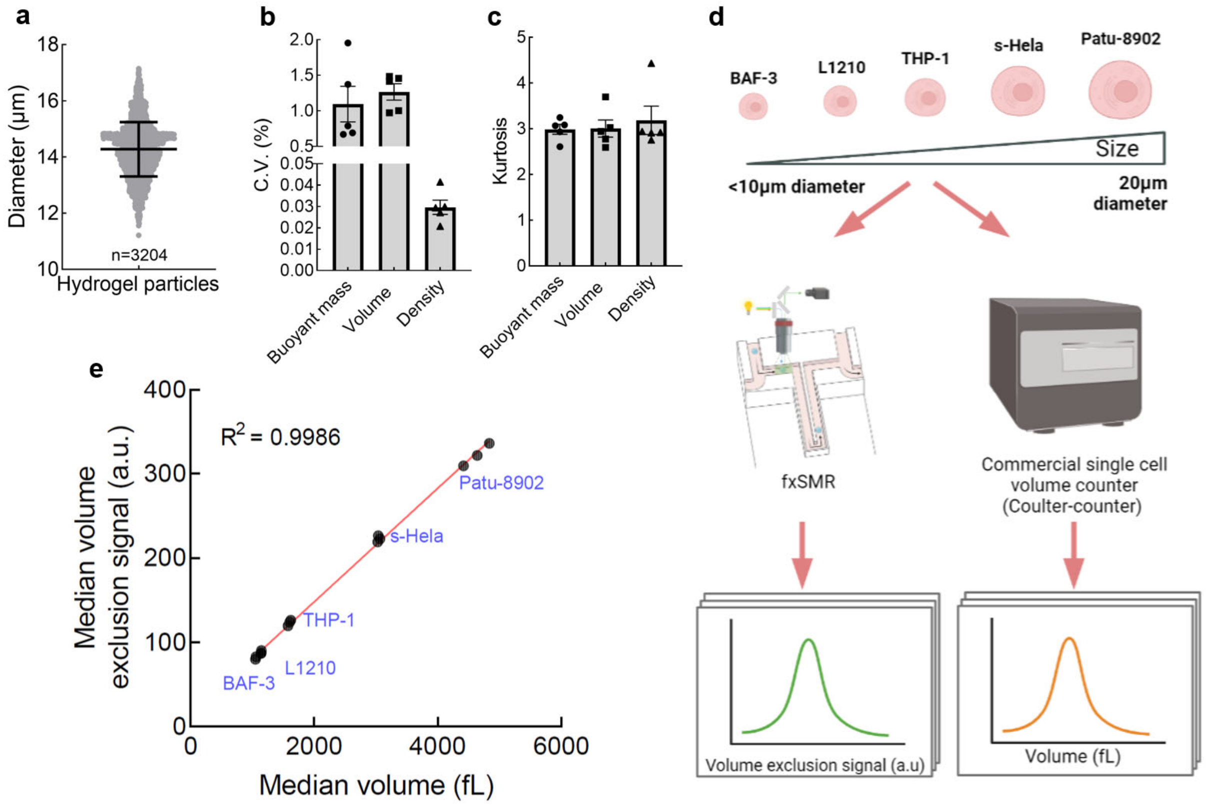


**Supplementary Fig. 1** | **Characterization of system accuracy and precision. a,** Size distribution of hydrogel particles used to characterize system precision. **b,** C.V. of buoyant mass, volume and density from single hydrogel particle trapping experiments, where the same particle was measured repeatedly (n = 5 independent experiments). **c,** Kurtosis of data from (b); Expected kurtosis of a normal distribution is 3. **d,** Workflow on characterizing the accuracy of volume measurements from fluorescence exclusion by comparing fxSMR results to coulter-counter measurements. **e,** Median volume of each cell line (n = 14 samples in total) from fxSMR (y-axis) vs coulter-counter (x-axis). y-intercept = 13.11 a.u.

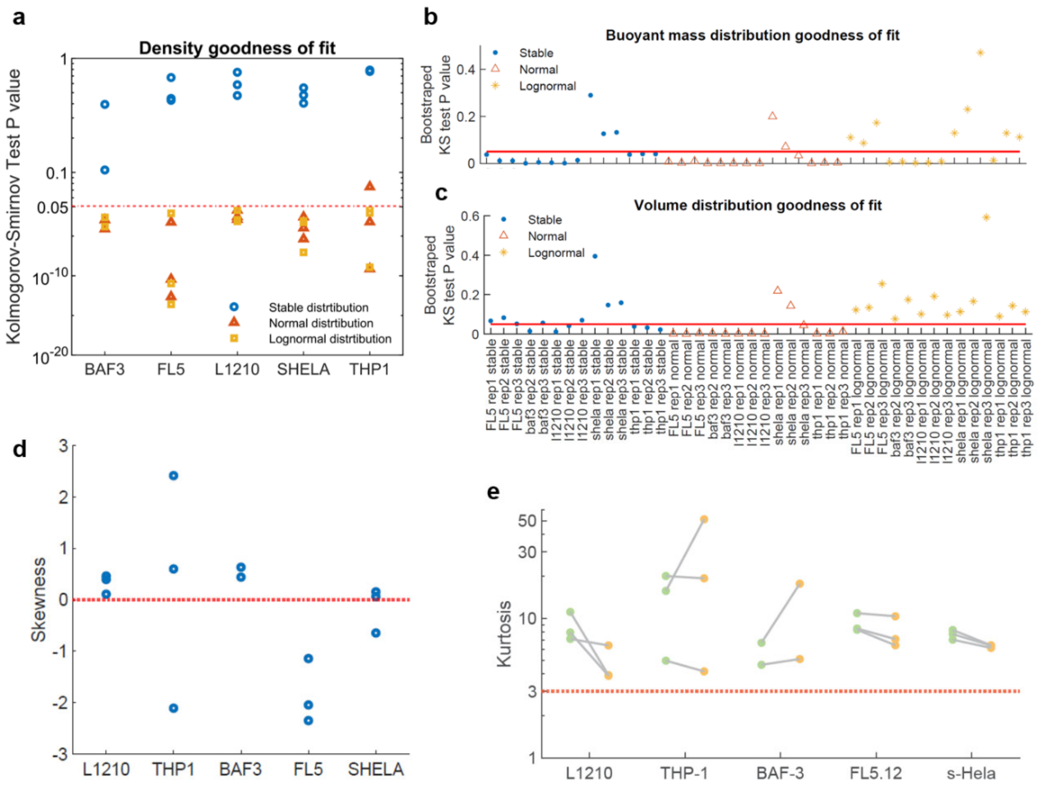


**Supplementary Fig. 2 | Characterization of density mass and volume distributions of proliferating mammalian cell lines. a,** Kolmogorov–Smirnov test on the mean goodness of fit on bootstrapped density distribution, when using stable, normal, and lognormal distribution fits; A p-value > 0.05 means the distribution is well-fitted. **b,** Kolmogorov–Smirnov test on the mean goodness of fit on bootstrapped buoyant mass distribution; Red bar denotes p-value of 0.05. **c,** Kolmogorov–Smirnov test on the mean goodness of fit on bootstrapped volume distribution; Red bar denotes p-value of 0.05. **d,** Skewness of each cell line when excluding 1% of outliers. **e,** Kurtosis of cells with light buoyant mass (lower 50% of the population, in green) and heavier buoyant mass (upper 50% of the population, in yellow) for each cell line. For (a-e), n = 3 biological replicates for FL5.12, L1210, s-Hela, THP-1; n = 2 biological replicates for BaF3.

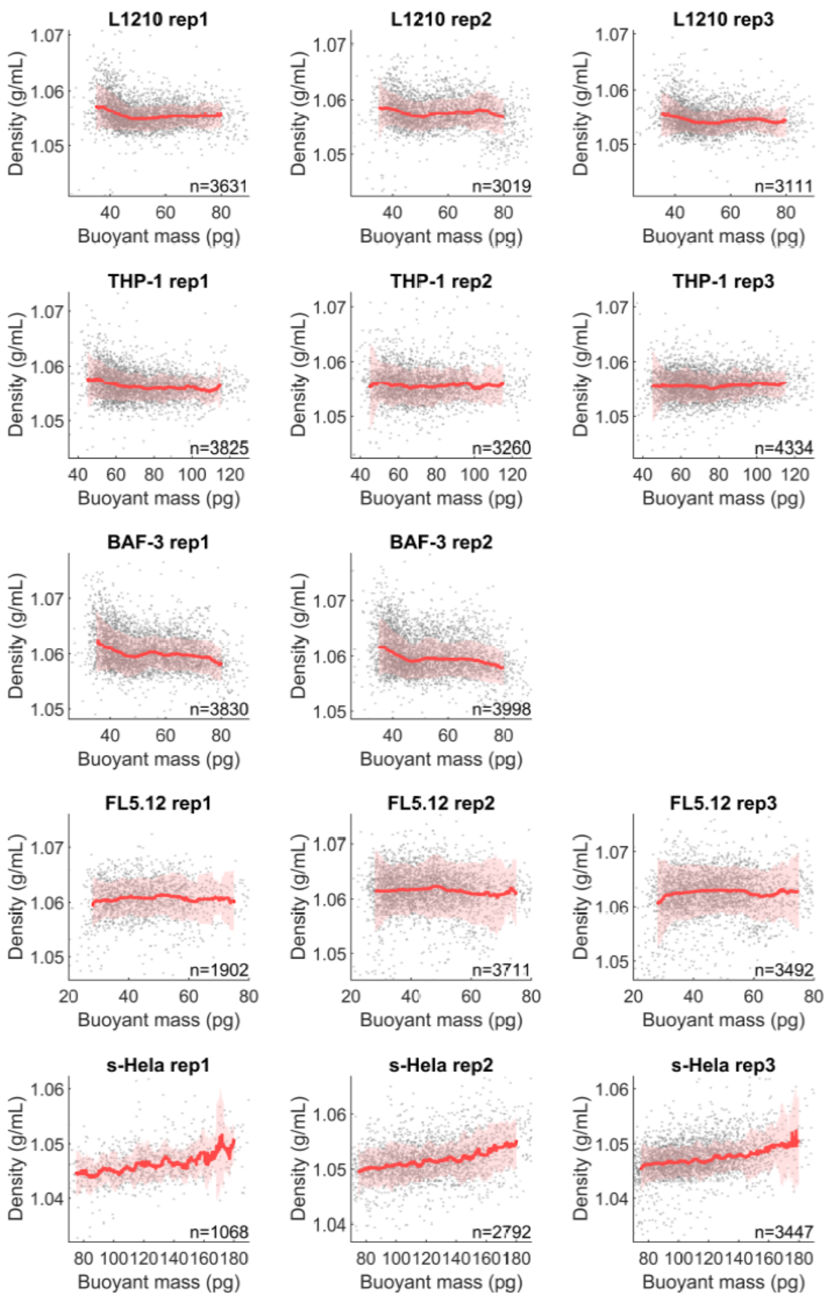


**Supplementary Fig. 3 | Density vs buoyant mass of proliferating mammalian cell lines.** Each grey point indicates a single cell; Red lines show moving median density as function of buoyant mass; The filter window is ± 5 pg and the step size is 0.1 pg; Shaded red regions indicate ± standard deviation of density in the filtered population; n values refer to the number of individual cells.

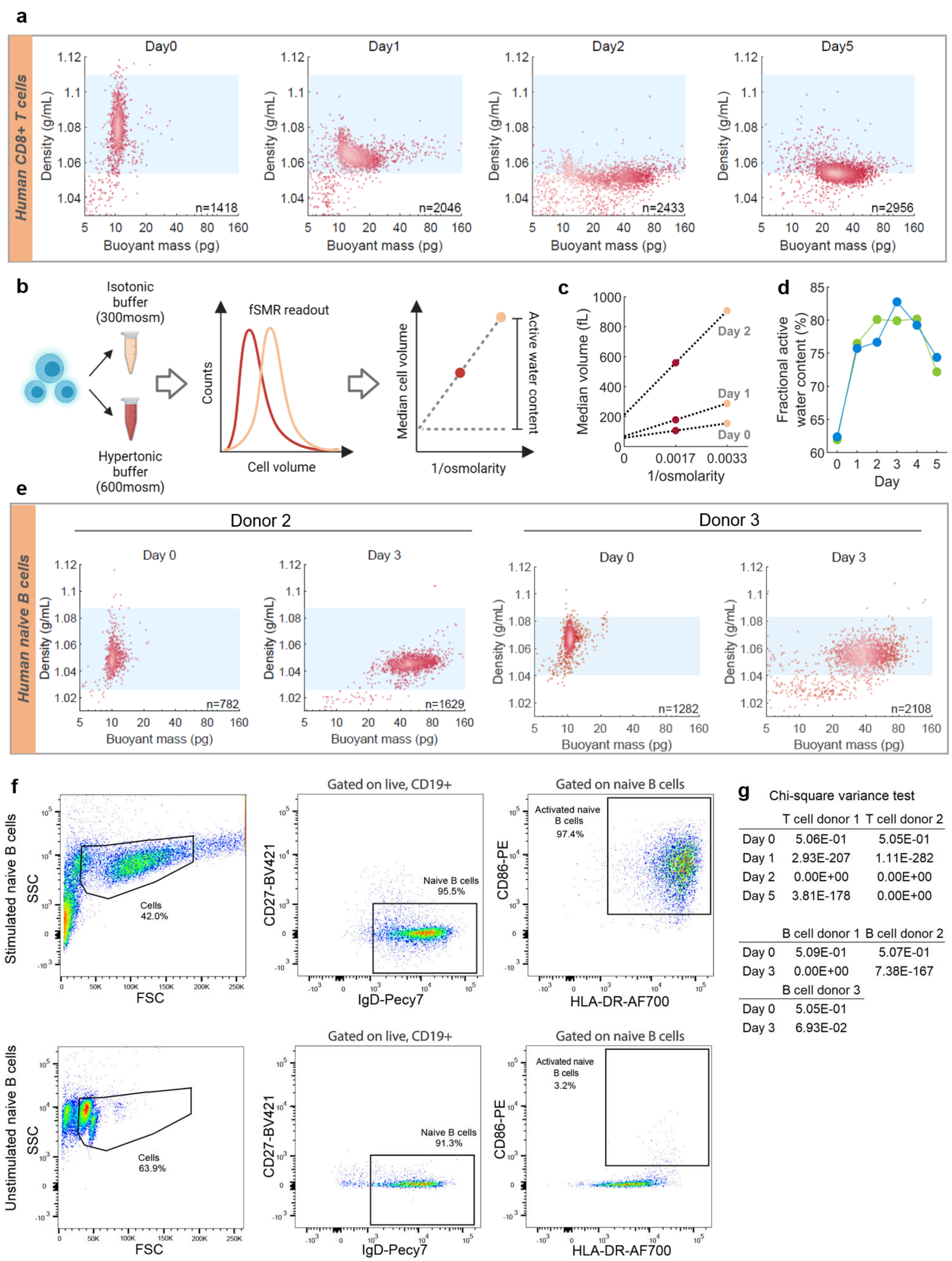


| | T cell donor 1 | T cell donor 2 |
|---|---|---|
| Day 0 | 5.06E-01 | 5.05E-01 |
| Day 1 | 2.93E-207 | 1.11E-282 |
| Day 2 | 0.00E+00 | 0.00E+00 |
| Day 5 | 3.81E-178 | 0.00E+00 |

| | B cell donor 1 | B cell donor 2 |
|---|---|---|
| Day 0 | 5.09E-01 | 5.07E-01 |
| Day 3 | 0.00E+00 | 7.38E-167 |
| | **B cell donor 3** | |
| Day 0 | 5.05E-01 | |
| Day 3 | 6.93E-02 | |

**Supplementary Fig. 4 | Density dynamics of human lymphocytes during transition between quiescence and proliferation. a.** Scatter plots of mass vs density showing the T cell dynamics post activation for donor 2; Blue areas indicate the density range of quiescent T cells at day 0, with upper and lower bounds indicate the 1st and 99th percentile of the density distribution; C.V. of density at day 0, 1, 2 and 5 are 1.035%, 0.492%, 0.436%, 0.445% accordingly; n values refer to the number of individual cells. **b,** Schematic showing the measurement of average active water content using Van't Hoff's osmosis principle; Each T cell sample is separately measured in an isotonic buffer (peach color) and a hypertonic buffer (dark red color); Dotted line connecting the median cell volumes indicates a linear fitting and the intercept at the y-axis indicates inactive volume (total volume minus the active water content). **c,** Representative median cell volume of one donor at days 0, 1, and 2 post activation, measured at two osmolarities; Dotted lines indicate linear fitting results. **d,** Fractional active water content (active water content / total cell volume) for T cells from both donors as a function of time. **e,** Representative scatter plots of mass vs density showing B cells at day 0 and 3 post activation for samples obtained from donor 2 and 3; Blue areas indicate the density range of quiescent B cells similar to (**a**). C.V. of density at day 0 and 3 are 0.998% and 0.495% for donor 2, 0.798% and 0.737% for donor 3 accordingly; n values refer to the number of individual cells. **f.** Representative flow cytometry plots showing the gating strategy for CD86+ activated naïve B cells population. The immunophenotype of stimulated (top panels) or unstimulated (bottom panels) naïve B cells after 3 days of cell culture is shown. Numbers adjacent to the gates represent cell frequencies. **g.** Chi-square variance test on T cell and B cell density during activation. Each number indicates the p-value when comparing the variance of density distribution from each condition to the variance of the day 0 population of the same donor. The null hypothesis is that two distributions have the same variance, and the alternative hypothesis is that the sample distribution has a smaller variance than the day 0 condition. The p-value of 0 refers to a p-value that is lower than what our calculation software (MATLAB) can determine.

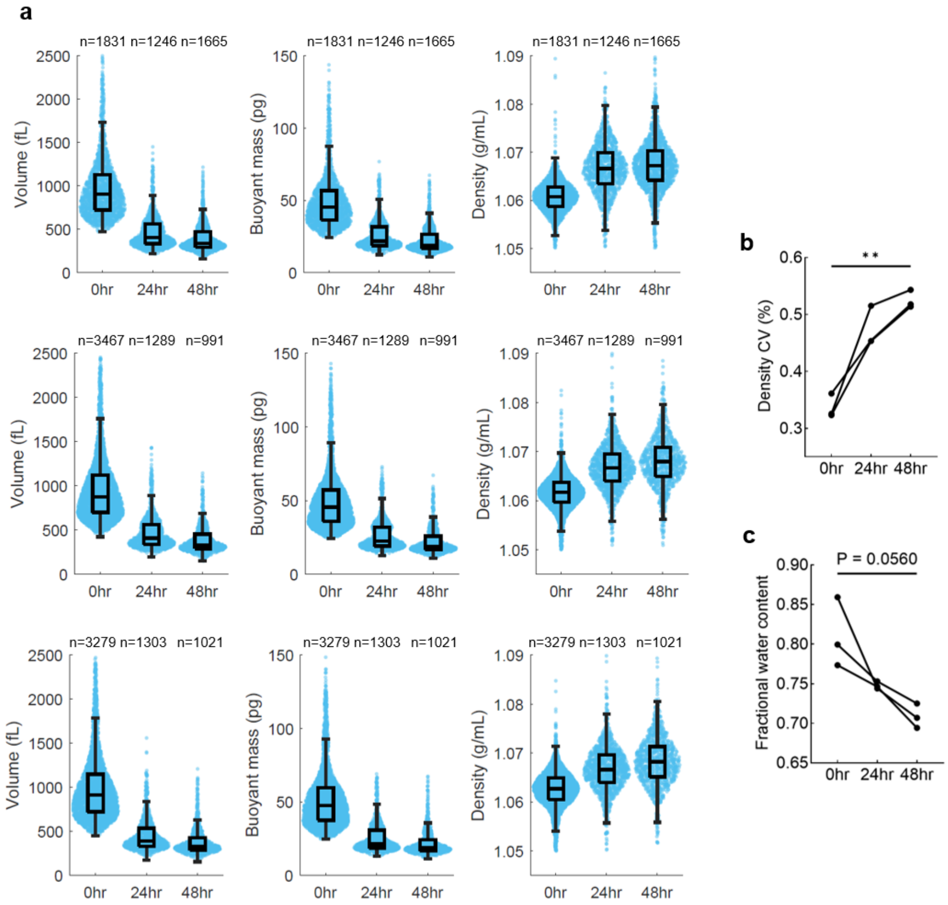


**Supplementary Fig. 5 | Density dynamics of murine pro-B lymphocytic cell line FL5.12 during cell cycle exit induced by growth factor depletion. a,** Box and whisker plots of FL5.12 cell volume, buoyant mass and density vs time after removing IL-3 supplement from the culture media; Each row shows a replicate condition; n values refer to the number of individual cells. **b,** Density C.V. vs time after IL-3 depletion; p-value = 0.0087 from two-tailed paired parametric t test between 0 and 48 hours; n = 3 biological replicates. **c,** Fractional active water content (active water content/total cell volume) vs time after IL-3 depletion; p-value is derived from two-tailed paired parametric t test between 0 and 48 hours; n = 3 biological replicates.

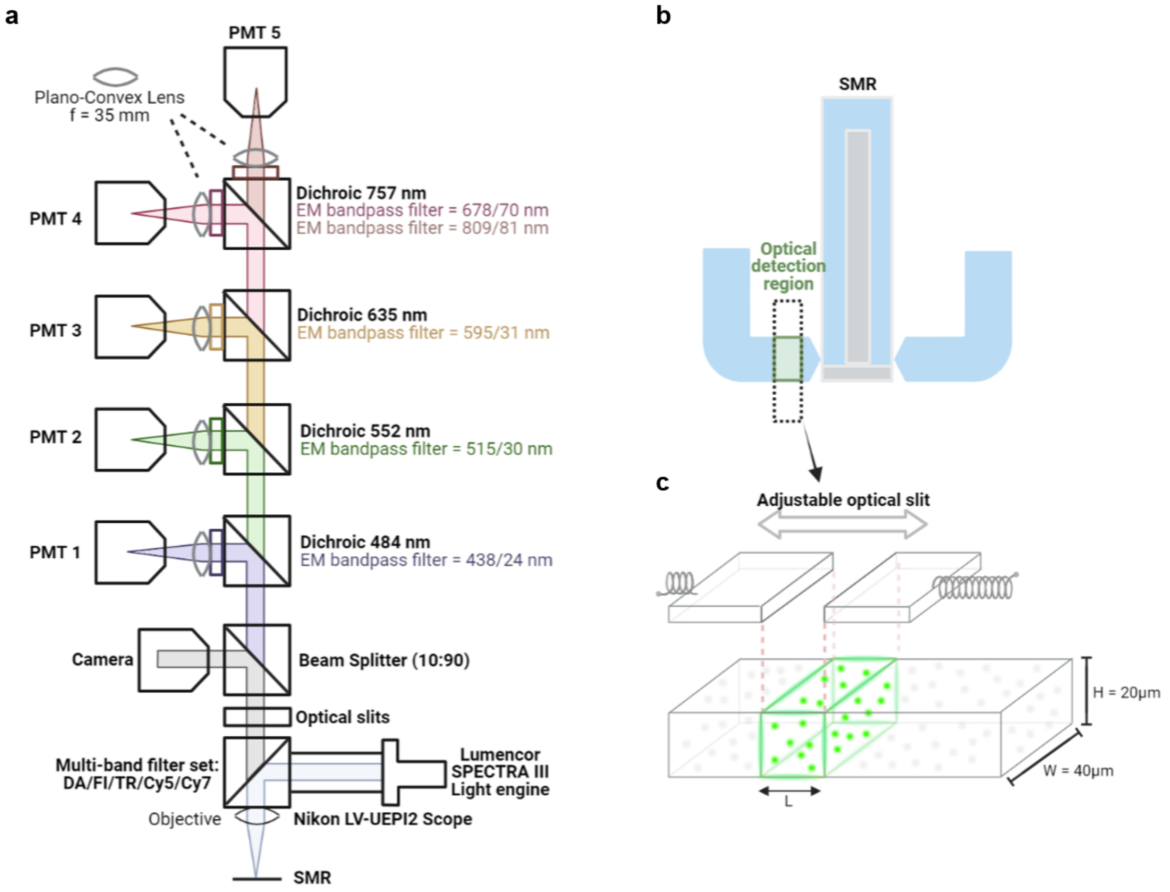


**Supplementary Fig. 6 | Optical setup for fluorescence exclusion-based volume measurements of cells in flow. a,** Schematics of an optical setup enabling multi-band fluorescence intensity measurements; Each colored light indicates light within a specific wavelength range; FITC-dextran fluorescence exclusion measurements are detected by PMT 2. **b,** Schematic of optical detection region where emission light is collected from, in reference to the SMR cantilever. **c,** Illustration of the optical detection region (green); Fluorescence exclusion baseline intensity is determined by the volume of the detection region, which depends on channel height (H), width (W), and the distance set by an adjustable optical slit (L).

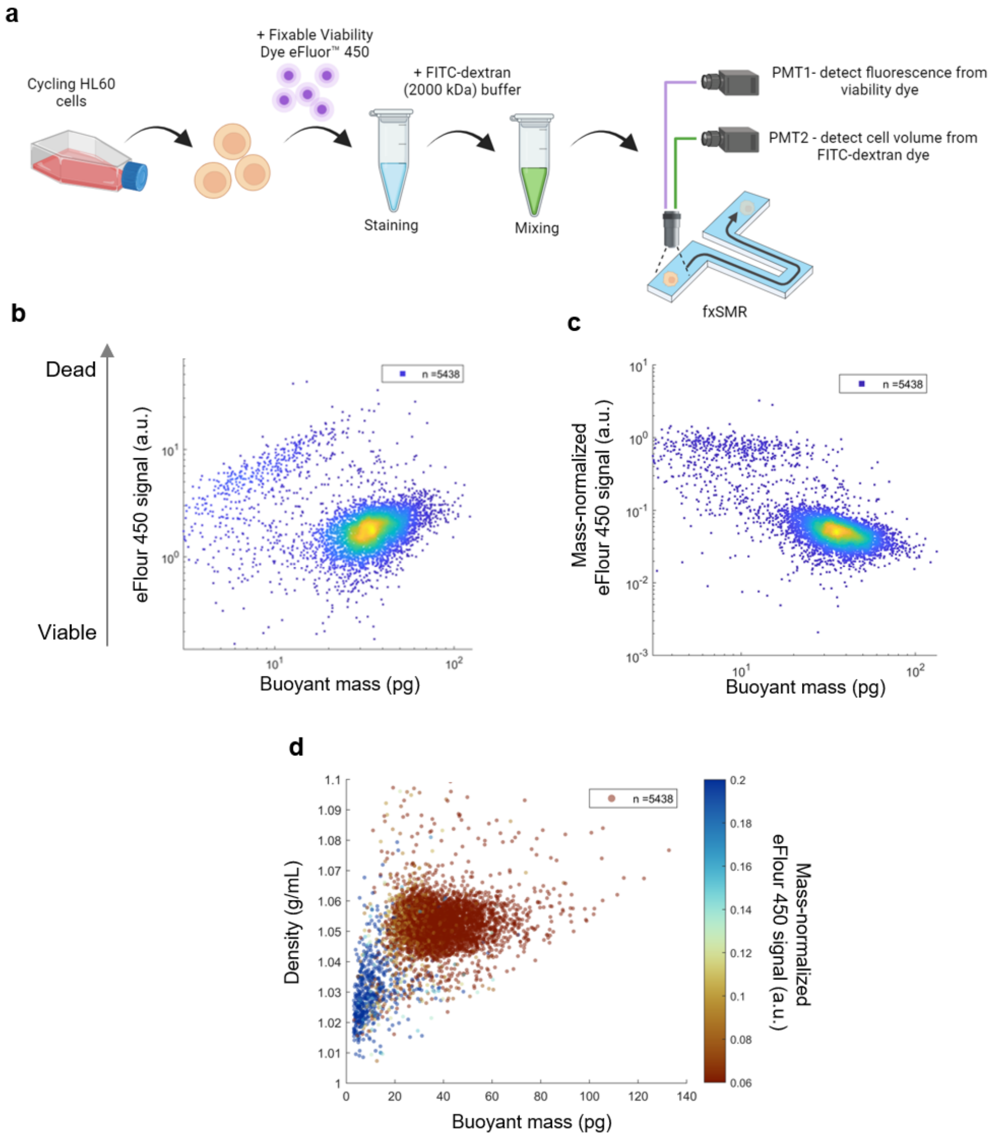


**Supplementary Fig. 7 | fxSMR resolves distinct clusters of viable and dead cells. a,** Schematic on experimental design for viability labeling of cycling HL60 cells and downstream fxSMR measurements. **b,** Scatter plot of viability vs buoyant mass on a population of HL60 cells; n value refers to the number of individual particles. **c,** Scatter plot of mass-normalized viability vs buoyant mass on the same population. **d,** Scatter plot of density and buoyant mass on the same population of cells; Color projection indicates the value of mass-normalized viability labeling; Dark red cluster indicates live cells and dark blue cluster indicates dead cells and debris.

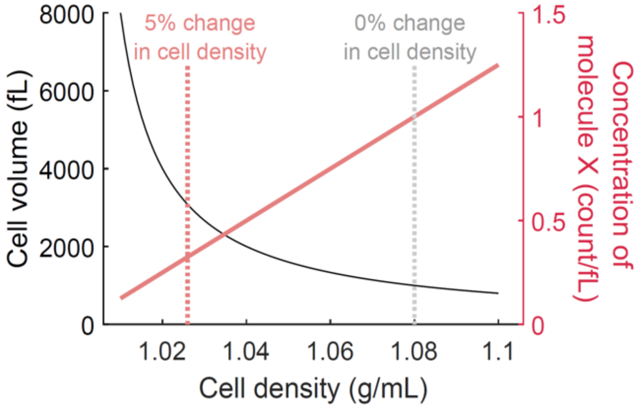


**Supplementary Fig. 8 | Simulated relationship between changes in cell density and cell volume or molecular crowding level.** Dry volume is dry density of the cell are treated as constants (200 fL and 1.4 g/mL accordingly). Molecule X is a hypothetical molecule with a defined concentration of 1 molecule per fL when the cell density is at 1.08 g/mL. Concentration of molecule is calculated by total molecule number (a constant) divided by cell volume.

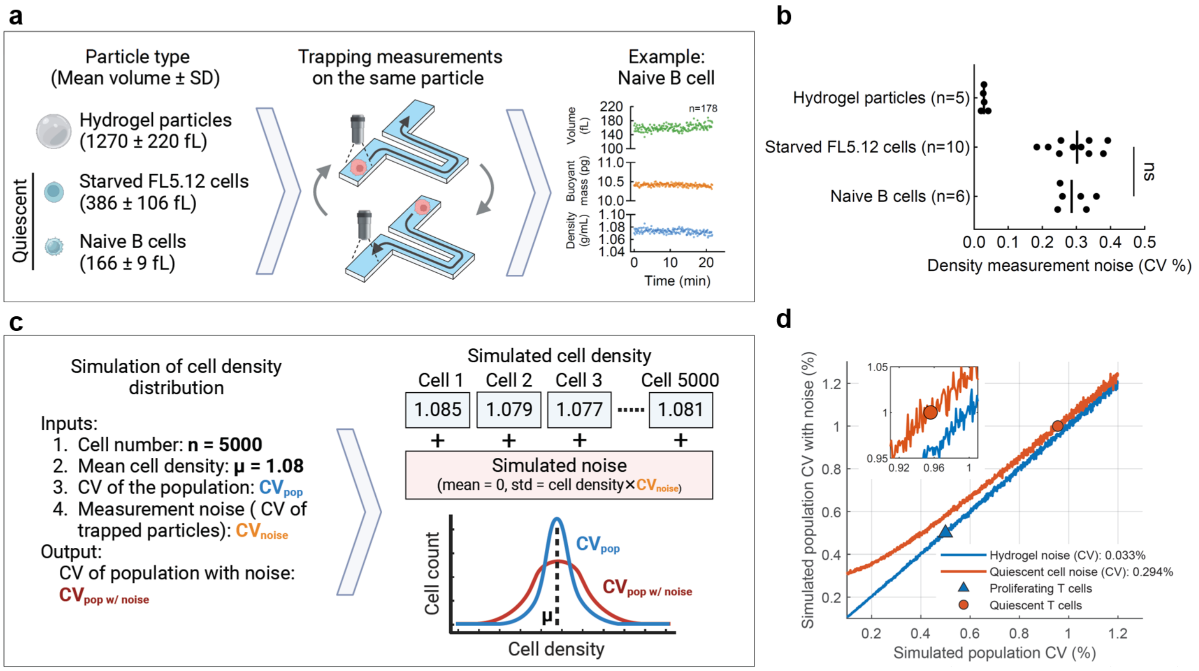


**Supplementary Fig. 9 | Characterization of density measurement noise for small particles and simulation on its effect on observed density heterogeneity in T cell populations. a,** Schematics on the experimental setup used to determine the precision of single cell density measurement for 3 types of particles (hydrogel particles, FL5.12 cells after 48 hours of IL-3 starvation, and naïve human B cells). The average volume of trapped particles and their standard deviations are reported in parentheses. **b,** Density measurement noise for each type of particle. Noise is defined by the C.V. of the density measurements from a single trapped particle. Each dot is a trapped particle. Unpaired parametric t-test between FL5.12 and B cells has a p-value = 0.934. **c,** Schematics on the simulation design to evaluate the effect of measurement noise on observed density heterogeneity of a cell population. **d,** Simulation results on input population C.V. ranging from 0.1% to 1.2%. The X axis represents the input variable, namely input 3 from (c). The Y axis represents the output variable, which is the observed population C.V. after accounting for the measurement noise. Line color indicates the simulation result when the noise term, namely input 4 from (c), is set to the average noise values reported in (b). Blue line indicates hydrogel particles, and red line indicates the quiescent cell types, which include both FL5.12 and B cell. Observed population density C.V. for proliferating and quiescent T cells from **Fig. 3** are overlaid on the line profiles, which represent the corresponding noise levels expected in each T population due to their difference in cell size.

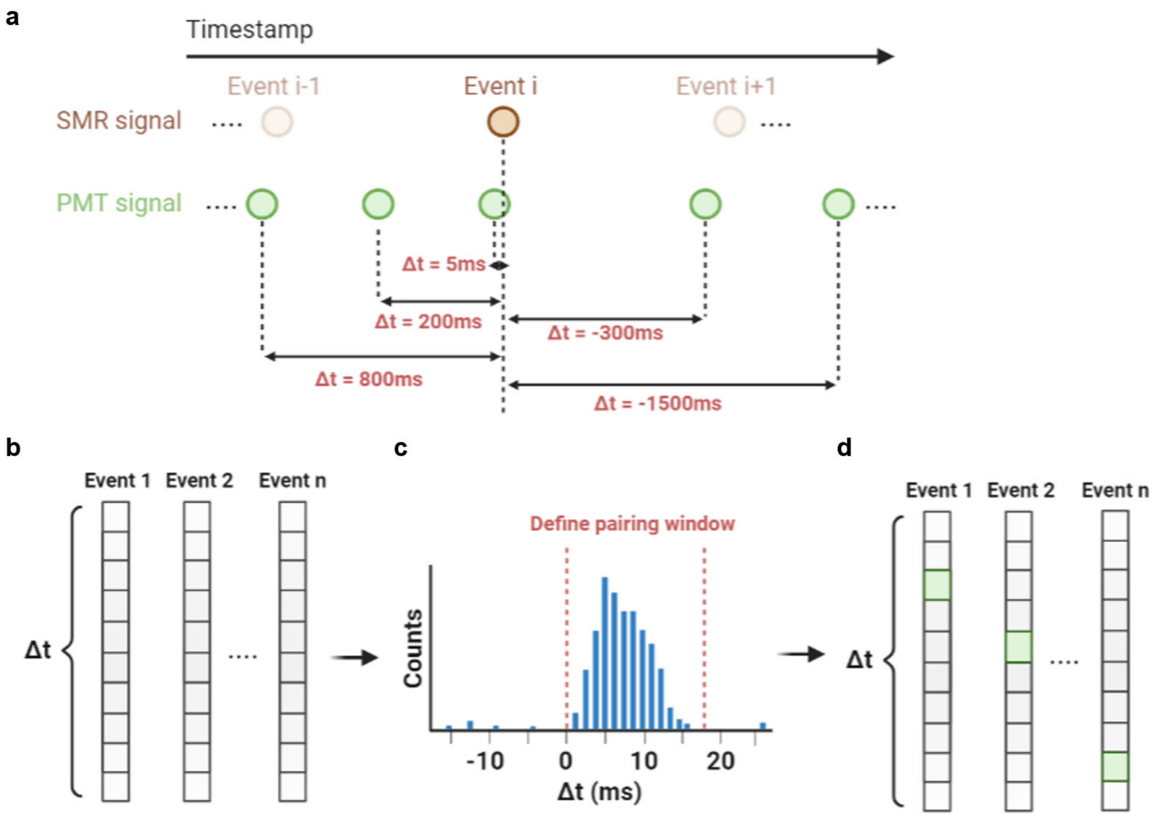


**Supplementary Fig. 10 | Design of SMR (mass) and PMT (volume) data pairing algorithm. a,** Schematic of temporal sequence for mass signals (brown) and volume signals (green) when aligned by their time stamps; Δt denotes the time difference between a SMR event i and a neighboring PMT signal. **b,** Schematic of the first step of the pairing algorithm: for each event in (**a**), an array of Δt is computed where each element is the Δt between this SMR event and a PMT signal; The length of the array is the total number of PMT signals measured in a sample. **c,** Schematic of the second step of pairing algorithm: a histogram of all the Δt values from all events in (**b**) is displayed, which results in a unimodal distribution of Δt; A user-defined range of Δt will be used for pairing. **d,** Schematic of the last step of the pairing algorithm: all SMR events that have exactly one Δt value that is within the user-defined range from (**c**) will be selected and paired to the corresponding PMT.

**Supplementary Note 1. Viable cell gating on fxSMR data.**

Viable cell classification for all single-cell measurements was done by manual gating using buoyant mass and density. This gating strategy was designed based on the assumption that cell buoyant mass and density could distinctly resolve live and dead cells. We validated this approach by first staining a population of HL60 cells with a standard viability dye that positively labels dead cells (ThermoFisher, 65-0863-14) and then performing a fxSMR measurement where mass, volume, density and viability dye emission level were captured for every cell (**Supplementary Fig. 7a**). The viability dye has an excitation/emission range of 405/450 nm, which does not overlap with the FITC-dextran used for measuring cell volume. Simultaneous detection of both fluorescence signals was achieved by the multi-band fluorescence detection setup of the fxSMR system (**Supplementary Fig. 6a**). Since the intensity of the viability label had a correlation with cell size, we performed a normalization step where the label intensity of each cell was divided by its buoyant mass, which largely removed the size-dependency of the viability label and can differentiate between live and dead cells at a threshold value of 0.2 a.u in mass-normalized viability signal (**Supplementary Fig. 7b,c**). When visualizing the mass-normalized viability signal together with cell mass and density, we observed that the clusters of viable and dead cells were distinctively resolved by cell mass in combination with cell density (**Supplementary Fig. 7d**).

**Supplementary Note 2. Relationship between cell density and molecular crowding.**

To understand the quantitative relationship between cell density and molecular crowding, we performed a simulation which shows that a small change in cell density (<5%) can reflect radical changes in the physicochemical milieu of the cytoplasm (**Supplementary Fig. 8**). In this simulation, cell density changes 5% purely due to water uptake, which indicates an approximately 2-fold increase in cell volume, thereby diluting all intracellular molecular components by ~2-fold. Such an intracellular dilution would influence, for example, phase transitions and enzymatic reaction rates.

A number of recent publications have shown that osmotically induced cell volume changes, which correspond to ≤ 5% cell density change, can radically alter phase transitions in a variety of mammalian models[1–4], thereby impacting processes involving nucleolus, processing bodies, stress granules, and other phase-separated organelles. For example, to achieve a 2-fold change in cell volume via osmotic shock, extracellular osmolarity has to change ~2-fold, according to Van Hoff's law. Thus, a 5% density change could have similar consequences as exposure of cells to 150 mOsm or 600 mOsm environments (assuming cells normally reside in a 300 mOsm environment). Studies on osmolarity-induced phase transitions have shown that even a small change in osmolarity (e.g. change of 50 mOsm, or 0.78% change in cell density) can alter phase transitions[1,2].

The 2-fold volume change due to water uptake can also impact enzymatic reaction rates. Let's imagine an enzymatic reaction where the substrate concentration is significantly below the $K_m$ of the reaction, as is the case for several enzymes in the central carbon metabolism[5]. According to Michaelis–Menten reaction kinetics, the initial reaction rate ($v_0$) for such cases can be estimated as follows:

$$v_0 \approx \frac{k_{cat}}{K_M}[E][S]\,, if\ [S] \ll K_M \qquad (1)$$

where [E] and [S] represent the enzyme and substrate concentrations, respectively. A 2-fold volume increase would decrease such enzymatic reaction rates ~4-fold. We note that this is a simplified example, which does not take into account the complex and crowded nature of the intracellular environment[6] and that not all enzymes operate in a substrate limited regime. Yet, these calculations highlight that a small (~5%) change in cell density can have major consequences for cell function and physiology.

**Supplementary Note 3. Effect of density measurement noise on observed density heterogeneity.**

To understand the effect of particle size on measurement uncertainty, we performed single-particle trapping experiments where we profiled small, quiescent cells including IL-3-starved FL5.12 cells and naive B cells (**Supplementary Fig. 9a**). These quiescent cells are ~5-10 times smaller than the hydrogel particles used to characterize system precision in **Fig. 1**, and they are in the size range of quiescent T cells. By repeatedly measuring the same cell within the device, we found that density measurements have an average C.V. of 0.294% (**Supplementary Fig. 9b**) compared to an average C.V. of 0.033% for the larger hydrogel particles. Although the C.V. increased, it is still considerably lower than the C.V. we measured for quiescent T cells of ~1% (**Fig. 3**), which indicates we are measuring inherent biological variability.

Since this noise value is higher than what was observed in the large hydrogel particles, we performed a simulation to more deeply understand the degree to which the measurement uncertainty affects the observed cell density heterogeneity in a population (**Supplementary Fig. 9c**). We first simulated the density values for a population of single cells (n=5000) that has a mean of 1.08 g/mL and a population C.V. (selected between 0.1%-1.2%, which covers the range of quiescent and proliferating T cells). We then added a noise term to each cell density value that represents the measurement uncertainty. The noise term follows a normal distribution with a C.V. defined by the uncertainty of either the hydrogel trapping data (a proxy for proliferating T cells) or the FL5.12 and B cell trapping data (a proxy for quiescent T cells). As shown in **Supplementary Fig. 9d**, the simulation reveals that when the cell population has low density variability (<0.4%), the observed density C.V. is more influenced by measurement noise. At higher density variability (>0.7%), the population C.V. with noise is roughly linear to the population C.V. without noise. This suggests that the measurement noise observed in the small cells has less influence when there is high intrinsic biological heterogeneity. After accounting for measurement uncertainty, the simulation shows that the 1% density C.V. that we measured in quiescent T cell populations translates to a population C.V. of ~0.96% (**Supplementary Fig. 9d**). This is higher than the ~0.5% C.V. observed in proliferating T cell density distributions.

**Supplementary Note 4. Data analysis pipeline.**

Raw data processing was carried out in MATLAB. Frequency peak analysis of raw SMR and PMT signals was processed as in previous studies[7,8]. For fluorescence exclusion volume measurements, PMT data were processed by a median filter and a moving-average filter (filter size = 5 data points for both filters). Event identification was based on negative thresholding on the fluorescence level, as defined by a decrease in fluorescence that is larger than three times the standard deviation of the baseline. Volume signals were computed by dividing the absolute value of the fluorescence decrease by its surrounding baseline signals that are within 100 data points from the identified peak.

We then identify and remove low-confidence volume signals to account for several factors that affect measurement quality. These factors include flow rate variations, out-of-focus events as well as unequal distribution of dye molecules around the cell. While the intensity of fluorescence emission from the excited region mainly depends on its volume, alterations in flow rate may cause differences in intensity if the dye molecules are prone to photobleaching. Similarly, out-of-focus event may cause changes in emission intensity when the volume of the excited region remains unchanged. Both factors reduce the quality of the detected signals but can be identified by a change in the baseline fluorescence intensity level. A change in flow rate will create a slope in the baseline, and out-of-focus will result in a change in average baseline intensity. In addition, signal quality can be reduced by a change in baseline fluorescence levels before and after cell passing, because dye molecules can accumulate near the cell as it passes through a constriction[9]. This bias can be identified by a significant change in baseline intensity after cell passing.

Low-confidence volume signals are identified and removed from the output given a predefined list of criteria: (1) a slope in baseline with an absolute value higher than 0.002 V/point; (2) baseline values that substantially differ from the median baseline value of the first 30% of identified peaks (±10% of median baseline amplitude); (3) signals that has a substantial difference between the left and right-side signal baselines (larger than 5% of the peak amplitude).

Data pairing between SMR and PMT signals was carried out after independent peak identification of SMR and PMT signals (**Supplementary Fig. 10**). Each SMR or PMT event had a distinct time stamp in computer real-time collected in the LabView software. We start by aligning one directional array for SMR and PMT time stamps, where the length of the array was the number of identified events, and each element was the timestamp of one distinct event. We assumed that every event in the SMR array should have a matching event from the PMT, i.e. the mass signal and volume signal of the same cell, but with a short time delay because the PMT signal was acquired at a different location than the SMR signal (**Fig. 1a**). The time difference between the two can vary due to slightly different flow velocities from one cell to another. However, on a populational level, we expected the time difference to have unimodal distribution since the pressure settings were identical through each experiment. By identifying the range of this distribution, we were able to uniquely pair the SMR signals with the PMT signals. The pairing pipeline started with computing the time difference ($\Delta t$) between each SMR event and every PMT event, which resulted in $m \times n$ number of $\Delta t$, where m and n are the total numbers of SMR and PMT events, respectively. A histogram of all $\Delta t$ values was then generated and the distribution typically centered between -10 and +10 ms depending on the exact position of the optical detection region in comparison to the SMR cantilever. Because the average time between consecutive cells was around 120 ms, there is a very low likelihood that two subsequent signals appear within ±10 ms. A manual selection of pairing range on the $\Delta t$ histogram was required due to sample-to-sample variability in flow rate. Then SMR and PMT data were paired if there was a unique one-to-one matching within the user-defined Δt pairing window. Doublet events, where there is more than one PMT or SMR signal in the same Δt window, were excluded. The pairing algorithm typically yielded a pairing rate of ~90% for the SMR signals in each measurement.
